# Convergent evolution of genes controlling mitonuclear balance in annual fishes

**DOI:** 10.1101/055780

**Authors:** Arne Sahm, Martin Bens, Matthias Platzer, Alessandro Cellerino

## Abstract

Complexes of the respiratory chain are formed in a complex process where nuclearly-and mitochondrially-encoded components are assembled and inserted into the inner mitochondrial membrane. The coordination of this process is named mitonuclear balance and experimental manipulations of mitonuclear balance can increase longevity of laboratory species.

Here, we investigated the pattern of positive selection in annual (i.e. short-lived)and non-annual (i.e. long-lived) African killifishes to identify a genomic substrate for evolution of annual life history (and reduced lifespan).

We identified genes under positive selection in all mitonuclear balance: mitochondrial (mt) DNA replication, transcription from mt promoters, processing and stabilization of mt RNAs, mt translation, assembly of respiratory chain complexes and electron transport chain. Signs of convergent evolution are observed in four out of five steps. This strongly indicates that these genes are preferential genetic targets for the evolution of short lifespan and annual life cycle

## Introduction

It is well-established that single-gene mutations can increase longevity in model organisms (Gems and Partridge 2013). One conserved mechanism that recently emerged is mitonuclear balance. Mitonuclear balance describes the process that coordinates the synthesis and assembly of mitochondrial respiratory chain complexes comprising mitochondrially-and nuclearly-encoded components (Dillin, et al. 2002; Lee, et al. 2003; Copeland, et al.2009; Houtkooper, et al. 2013; Baumgart, et al. 2016). The genetic basis of lifespan evolution is a less investigated topic, partly due to scarcity of model taxa with divergent lifespans and genomic resources. Killifishes of the genus Nothobranchius are adapted to an annual life cycle that is dictated by the periodicity of the monsoon and inhabit seasonal savannah ponds in Eastern Africa whose duration limits the lifespan of the adults to less than one year. Embryos survive the dry season, encased in the dry mud, in a state of developmental arrest (diapause) (Cellerino, et al. 2015) that is characterized by profound changes in mitochondrial (mt) physiology (Podrabsky and Hand 1999; Duerr and Podrabsky 2010). Variations in the aridity of the habitats are associated to parallel evolution of lifespan in two Nothobranchius lineages (Terzibasi Tozzini, et al. 2013). These fish can be cultured in captivity and have become an experimental model system to investigate the genetic basis of the evolution of life-history traits, diapause and aging (Cellerino, et al. 2015). The genome of the species *N. furzeri* was recently sequenced and this allows to search globally for genes under positive selection (Reichwald, et al. 2015; Valenzano, et al. 2015).

## Results and discussion

We performed genome-wide scans for positive selection based on recently assembled transcriptomes of six species of the annual genus Nothobranchius and of *Aphyosemion striatum* as a representative species of the closest non-annual genus (Reichwald, et al. 2015). Our aim was to identify genes potentially involved in the evolution of annual life history (and reduced lifespan) (Fig.1).

**Fig. 1.**
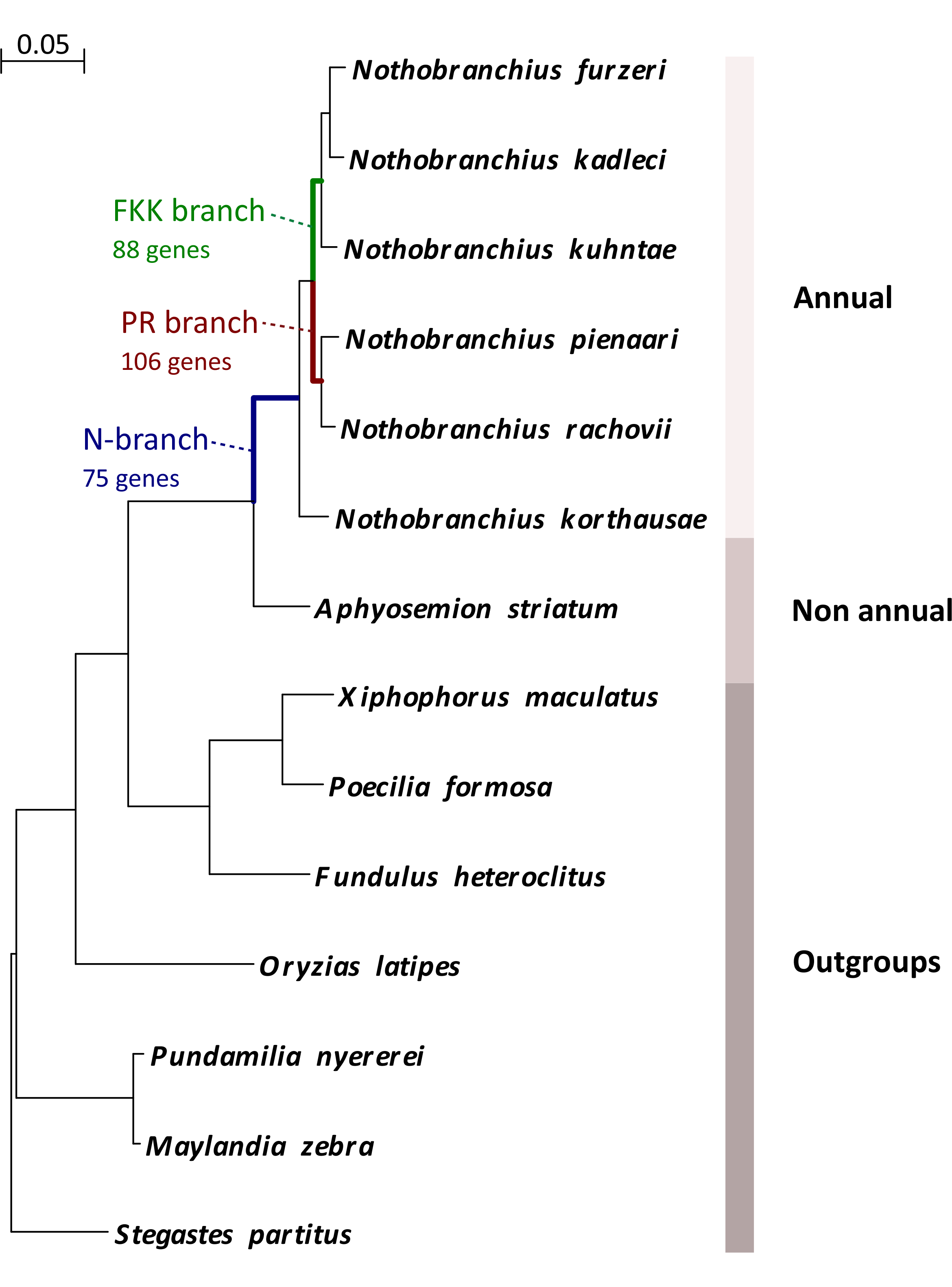
Phylogeny of the analyzed species and their life history Maximum-likelihood nucleotide-based phylogenetic tree of species that were used for genome-scale scans for positively selected genes. Outgoups from Ovalentaria are indicated as well as the three branches that are reported in the text. The alignment is based on concatenation of 4865 genes. The represented tree is the consensus of 1046 different trees created by splitting the alignment in fragments of 15 knt and calculating a tree for each fragment. The calibration bar refers to substituions per nucleotidic site

### Branch selection

We previously found seven genes under positive selection in *N. furzeri,* the shortest-lived species of the genus, and one in *N. pienaari,* another very-short-lived species, using the other six species of Nothobranchiidae as outgroups (Reichwald, et al. 2015). Here, we obtained from GenBank the RefSeq mRNA sequences of the phylogenetically closest outgroups from Ovalentaria (Fig 1) and analyzed the pattern of positive selection along three internal branches of the tree (Table S1). To identify events of convergent evolution, i.e. genes or functional categories that were positively selected in more than one branch, we selected the branches leading to the last common ancestors (LCAs) of *N. pienaari* and *N.rachovii* (PR-branch) and *N. furzeri, N. kadleci* and *N. kuhntae* (FKK-branch). These two branches diverged in the Pleistocene, share the same distribution and species belonging to the two clades can be found sympatric in the same pond (Dorn, et al. 2014). They represent therefore independent adaptations to the paleoclimatic changes of that period that was characterized by long-term progressive aridification of East Africa (Dorn, et al. 2014) and likely they were both subject to continued selection on adaptations linked to annual life cycle.

### Respiratory complex

In the branch of the LCA of the annual species (N-branch), we found 75 genes under positive selection (p<0.05,branch-site test) and among these, four code for components of the mitochondrial respiratory chain complex I (GO:0030964, fold-enrichment=14, p=0.0002, Fisher's exact test). Therefore, evolution of annual life cycle is coincident with strong positive selection on mitochondrial respiration. This is in line with the evidence that diapause is linked to profound remodeling of mitochondrial physiology (Podrabsky and Hand 1999; Duerr and Podrabsky 2010). Three further genes of complex I are under positive selection in the PR-branch,(fold-enrichment=8.8, p=0.005, Fisher's exact test)and one further gene in the FKK-branch, indicating continued and convergent positive selection on complex I during the evolutionary history of Nothobranchius. No evidence was found for positive selection on the mitochondrially-encoded components of complex I.

### Mitochondrial biogenesis

Strikingly, nine genes were under positive selection in both the PR-and FKK-branches (Table S1). Among these, *areTFB2M* (transcription factor B2, mitochondrial) and POLRMT(polymerase (RNA) mitochondrial) that together with *TFAM* (transcription factor A, mitochondrial) form the ternary complex that transcribes the entire mitochondrial genome (Litonin, et al. 2010) and *FASTKD5* (fast kinase domain 5) that is necessary for processing of mitochondrial mRNAs (Antonicka and Shoubridge 2015). Further signs of convergent positive selection were evident at the level of functional gene groups. In addition to *FASTKD5; FASTKD1* and *LRPPPC* (leucine rich pentatricopeptide repeat containing),that control stability of mitochondrial RNAs (Sasarman, et al. 2010), were positively selected in PR-branch. Three proteins annotated as mitochondrial ribosome (MRPs) were under positive selection in each of the two branches (GO:0005761, fold enrichment=9.1 and 14.7,respectively, p=0.02 and 0.01, Fisher'sexact test for the-and FKK-branch, respectively). In addition, two recently identified MRPs(Koc, et al. 2013), *PTCD3* (Pentatricopeptide repeat-containing protein 3)and *CHCHD1* (Coiled-coil-helix-coiled-coil-helix domain containing protein 1) were positively selected in FKK-branch. Two further genes important for translation of mitochondrial RNAs were also positively selected: *MTIF3* (mitochondrial translation initiation factor 3) in FKK-branch and *TACO1* (Translational Activator of Mitochondrially Encoded Cytochrome C Oxidase I) in PR-branch. Respiratory chain complexes are large proteincomplexes that undergo multi-step assembly where nuclearly-and mitochondrially-encoded components are combined and inserted into the mitochondrial inner membrane(Ghezzi and Zeviani 2012).Several genes involved in this process were positively selected: *COX18* (COX18 Cytochrome C Oxidase Assembly Factor) (Sacconi, et al. 2009) in FKK-branch, *OXA1L* (oxidase (cytochrome c) assembly 1-like) (Stiburek, et al. 2007; Haque, et al. 2010) in PR-branch, *FOXRED1* (FAD-dependent oxidoreductase domain containing 1) (Fassone, et al. 2010) and *LYRM7* (LYR motif containing 7) (Sanchez, et al. 2013) inN-branch. Therefore, proteins necessary for expression of mitochondrially-encoded genes and assembly of respiratory chain complexes show signs of convergent evolution (Figure 2).Two further genes under convergent positive selection are possibly also involved in this process: *ETAA1*(Ewing's tumor antigen A1) and *APOA1* (apolipoprotein A1). Gene coexpression network analysis in *N. furzeri* indeed revealed that *ETAA1* and *APOA1BP*(apolipoprotein A1 binding protein) are co-regulated with MRPs and complex I components (Baumgart, et al. 2016).Finally, also one gene important for mtDNA replication, *SSBP1* (Single strand DNA Binding Protein 1, mitochondrial) (Korhonen, et al. 2004), was positively selected in the FKK branch.

**Figure 2.**
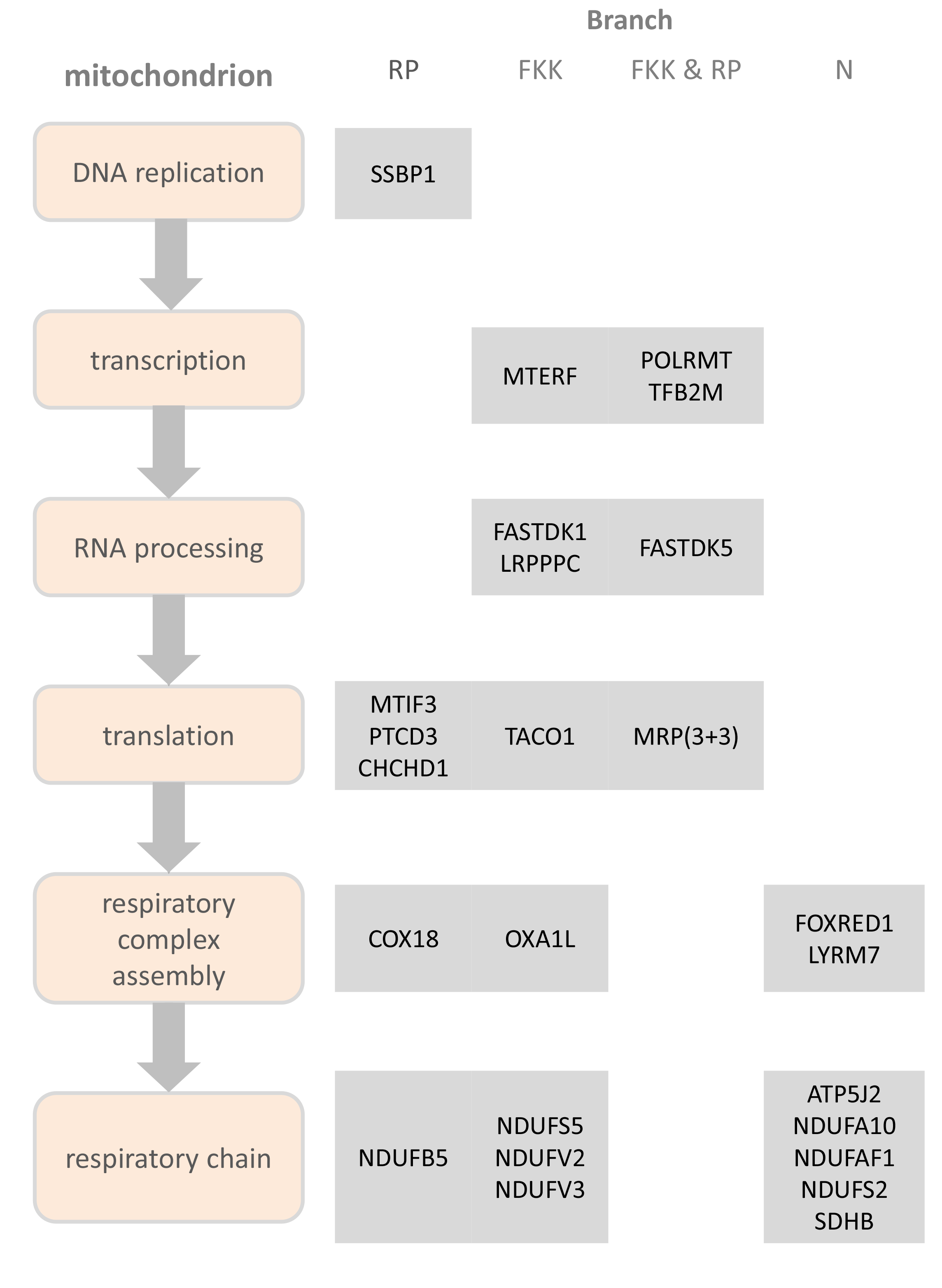
Convergent selection on genes controlling mitonuclear balance The genes under positive selection in each branch that are involved in the following processes: mtDNA replication, transcription from mitochondrial promoters, processing and stabilization of mitochondrial RNAs, translation (mitochondrial ribosome proteins selected: MRPL53, MRPS31 and MPRS26 in FKK-branch and MRPL23, MRPL3 and MTG2 in the RP-branch, respectively), assembly of respiratory chain complexes and electron transport chain are indicated for each of the three branches shown in Fig.1.

## Conclusion

The coordinated synthesis and assembly of mitochondrially-and nuclearly-encoded components of the respiratory chain (mitonuclear balance) is a conserved longevity mechanism that is controlled by MRPs (Dillin, et al. 2002; Lee, et al. 2003; Copeland, et al. 2009; Houtkooper, et al. 2013). In a recent longitudinal RNA-seq study in *N. furzeri,* we detected mitochondrial ribosomal proteins (MRPs) and complex I genes as central nodes in a network of genes negatively correlated with individual lifespan. Lower expression of these genes, as well as of *ETAA1* and *APOA1BP,* in young adult stage is associated with longer lifespans and partial inhibition of complex I prolongs lifespan in *N. furzeri* and partially reverts age-dependent transcriptome regulation in *N. furzeri* and zebrafish(Baumgart, et al. 2016). Here, we show that these genes underwent convergent evolution in annual fish, strongly suggesting that the control of mitonuclear balance represents a preferential genetic target for evolution of lifespans and life-history traits and are causally linked to evolution of short lifespan and annual life cycle.

## Materials and Methods

We used coding sequences of six Nothobranchius species (*N. furzeri, N. kadleci, N. kuhntae, N. pienaari, N.rachovii* N. korthausae) and *A. striatum* from transcriptome catalogs that were recently assembled by us (Reichwald, et al. 2015) with FRAMA (Bens, et al.). To increase the number of covered genes and isoforms we used for each species the union of two assemblies, one from a single end and one froma paired end sequencing approach. Codings sequences from seven additional outgroups *(Xiphophorus maculatus, Poecilia formosa, Fundulus heteroclitus, Maylandia zebra, Pundamilia nyererei, Stegastes partitus, Oryzias latipes)* were obtained from NCBI RefSeq (14.12.15) and assigned to ortholog groups by best-bidirectional blast against *N. furzeri.* Then, for each *N. furzeri* isoform the most similar isoform of each other species were determined by pairwise comparison. These sequences were required to have additionally at least a similarity of 70% with *N. furzeri* and 50% with each other species on protein level. The selected isoforms in each ortholog group were aligned with PRANK (Loytynoja and Goldman 2008), which is the alignment software of choice for positive selection analysis (Fletcher and Yang 2010; Jordan and Goldman 2012). The alignments were stringently filtered with GBLOCKS (Castresana 2000; Talavera and Castresana 2007) to remove unreliable alignment columns. Then, for each alignment the branch-site test of positive selection (Yang and Nielsen 2002; Zhang, et al. 2005) was applied: The respectively tested branch (LCA, FKK or PR) was marked as ‘foreground’ and all other branches were marked as ‘background’. The program CODEML from the PAML (Yang 1997, 2007) package were called separately for models M2a_0_ (model=2, NSsites=2; fix_omega=1, omega=1) and M2a (model=2, NSsites=2; fix_omega=0, omega=1) as described in the PAML User Guide (http://abacus.gene.ucl.ac.uk/software/pamlDOC.pdf). To calculate ap value the *χ*^2^ distribution with one degree of freedom was used to compare the likelihoods of both models: p=*χ*^2^(2*(ln(likelihood(M2a))-ln(likelihood(M2a_0_))),1). For 11089, 12735 and 11479 genes p-values were calculated in the N, FKK and PR branch respectively. Sites under positive selection were inferred by the Bayes Empirical Bayes method (Yang, et al. 2005) provided by CODEML. Sites that were predicted in a two amino acid frame next to a block which was deleted by GBLOCKS were removed and the p-value of the alignment corrected accordingly. Since high rates of false positive were detected in some automated genome-scale scans for genes under positive selection in the past (Mallick, et al. 2009;Schneider, et al. 2009; Markova-Raina and Petrov 2011), we demanded our final candidates to fulfil further filter criteria. Briefly, candidates were removed that:(I) had less than four species in the alignment or not had at least one species from each child taxon and the sister taxon of the respectively tested branch, (II) absolutely or relatively to few columns of the alignment remained after GBLOCKS filtering, (III) absolutely or relatively to few codons of the *N. furzeri* sequence remained after GBLOCKS filtering, (IV) disproportional d_N_/d_S_ ratios (e.g. >=100 in foreground branch or >1 in background branch) were calculated by CODEML or (V) had an unreliably high fraction of inferred positively selected sites. Finally we inspected all candidates on the FKK and PR branch manually as well asample of those on the LCA branch and removed ten additional candidates (<55)in total.

The phylogenetic tree was calculated based on concatenated alignment of those 4865 genes that had at least one isoform alignment in which all species were represented. The final tree is the consensus of 1046 different trees created by splitting the alignment in fragments of 15 k nt and calculating a tree for each fragment with DNAML from the PHYLIP (Felsenstein 2005) package.

